# A human-like EEG signature of cognitive control in the domestic dog

**DOI:** 10.64898/2026.02.11.705327

**Authors:** Ivaylo Borislavov Iotchev, Anna Kis, Márta Gácsi

## Abstract

In humans, theta (θ) band activity (defined as 4-8 or 5-7 Hz), measured over the frontal midline of the scalp, is an important EEG correlate of voluntary and conscious self-control. Theta waves specifically reflect the workings of the frontal lobes, and can therefore be useful in distinguishing cortical from more ancient control mechanisms. In dogs, inhibitory self-control is extensively studied, but mostly through behavioural tests. Here, we present a first inquiry into a possible EEG correlate of cognitive control in the domestic dog, by comparing short (∼30-second-long) EEG recordings from two conditions: passive wakefulness (baseline) versus a delayed gratification challenge (test). Both were recorded alternating, under similar conditions for each dog within the same session. In total, we collected and analysed 226 short recordings from fourteen dogs. Within and across animals, we found an increase in activity (test > baseline) that resembles human cognitive theta in frequency range (5-7 Hz) and scalp localization. Our results are a first demonstration that frontal midline theta in awake dogs can peak under similar conditions to those in humans. These findings indicate that compliant behaviour in dogs is under prefrontal control.

## Introduction

‘Cognitive control’ refers to those mechanisms of the mind-brain (Eisma et al., 2021) by which behavior is made to align with one’s goals and plans. Cognitive control allows information to be prioritized, structured, or ignored. It underlies the willed and conscious selection (van Schie et al., 2022) and suppression (Yamanaka and Yamamoto, 2010) of action choices.

In humans, cognitive control functions are localized in the (pre)frontal lobes (Ball et al., 1999; Swick et al., 2008; Zavala et al., 2018). The anatomy of the dog brain includes analogues of these structures (Cook et al., 2016; Czeibert et al., 2019; Szabó et al., 2023), but when and how the animals’ behavior is under prefrontal control is not thoroughly researched. Most of the evidence in dogs is derived from behaviour (reviewed in Olsen (2018)), overwhelmingly from studies on inhibitory control (Bray et al., 2014; Brucks et al., 2017; Cook et al., 2016; Olsen, 2022), i.e. the ability to suppress impulses, and task-irrelevant, or wrong responses. To date, only one study has linked this capacity to a hemodynamic response in the dog’s frontal lobes (Cook et al., 2016), but there are at least three reasons to expand our search for the neural substrates of canine cognitive control.

First, the many-to-one mapping problem in comparative cognition (Taylor, Alex et al., 2022) highlights a general problem with observational data: most behaviours can be achieved through different mechanisms or strategies. Response inhibition is no exception to this rule. It has been shown, for instance, that the cerebellum can inhibit motor responses in the absence of an intact neocortex (Hesslow, 1994). Another subcortical structure hypothesized to play a crucial role, specifically in the regulation of dogs’ behaviours, is the subthalamic nucleus (Medenica et al., 2025). In humans, the contributions of the frontal lobes and subthalamic nucleus are not mutually exclusive (Zavala et al., 2018), however it is common to observe structures that operate both within a hierarchy and independently. Since only neocortical inhibition is linked to voluntary and cognitive control of motion (van Noordt et al., 2022; Yamanaka and Yamamoto, 2010), a closer examination of the neural correlates of response inhibition, especially when they distinguish cortical from subcortical mechanisms, can highlight the level of cognitive engagement during action suppression.

Second, the behavioural data to date suggest that within a dog, the successful use of inhibitory control varies across tasks and contexts (Bray et al., 2014; Brucks et al., 2017). To understand which situations prompt prefrontal control (as opposed to subcortical strategies), it is important to pair behavioural and neural measures in canine cognitive experiments. To date, the only study to do so (Cook et al., 2016) used a highly artificial task, a variation of the human Go/No Go paradigm, on dogs lying motionless in an MRI scanner. In contrast, situations closer to everyday challenges, such as being forbidden to eat food off the floor, have not been examined in dogs using neuroimaging.

Third, while fMRI can most directly and reliably demonstrate the localization of task-relevant brain activity, follow-up research is hindered by the strong sensitivity of this technique to motion artefacts and its high cost of implementation (Montgomery, 2023). While restricted motion is the ideal condition for all neuroimaging methods, EEG is somewhat more forgiving and has well-established cleaning protocols e.g. for eye-movement artefacts (Ronca et al., 2024). EEG is also implemented at a lower cost (Montgomery, 2023) and does not require the animals to undergo lengthy training before they can be measured (Kis et al., 2014). All of these facts make it more likely to establish an extended series of cognitive control studies in dogs, if EEG concomitants of prefrontal control are identified.

In humans, elevated power in the theta band, defined either as 4-8 Hz (Eisma et al., 2021; van Noordt et al., 2022), or more narrowly 5-7 Hz (Inanaga, 1998; Jadeja, 2021), is linked to various types of cognitive control (Eisma et al., 2021), with a well-established link to response inhibition in particular (van Noordt et al., 2022; Yamanaka and Yamamoto, 2010; Zavala et al., 2018). It is thereby crucial that only prefrontal theta is linked to response inhibition, whereas theta recorded from the subthalamic nucleus was instead associated with response initiation (Zavala et al., 2018). This suggests that theta observed during action suppression will indicate neocortical control of behavior more reliably than theta observed during (re)active states. Moreover, theta linked to higher cognitive functions is consistently found at the frontal midline of scalp EEG recordings in humans (Eisma et al., 2021; Inanaga, 1998; van Schie et al., 2022). This distinction may prove crucial in analysing canine EEG for analogues of cognitive theta. There are currently only two active midline electrodes in most dog EEG experiments (Kulgod et al., 2025), but they appear sufficient to replicate humanlike differences in topography. For instance, a higher number of high-frequency spindles can be counted during slow-wave sleep over the more posterior midline electrode (Iotchev et al., 2019), as also seen in humans (Jobert et al., 1992) and rats (Terrier and Gottesmann, 1978).

In the present study, we devised a delayed gratification challenge for dogs, which required response inhibition towards the target, since it was visible and smellable, and remained within reach during the delay period. We compared the EEG from this test condition with representative baseline recordings from within the same session and, therefore, within the same overall conditions (same room, same time of day, same humans present, i.e. owner and experimenter). Since the test condition was linked to a greater need for self-inhibition, we expected to see higher frontal midline theta during test recordings compared to the baseline.

## Methods

### Ethics

By Hungarian law, non-invasive and harm-free studies with animals do not count as animal experiments subject to explicit permissions (For a corresponding ethical statement by the Hungarian Scientific and Ethical Committee of Animal Experiments, see PE/EA/2019-5/2017). All owners provided written consent to participate with their dogs in the study. They were informed of the type of data we collect, how the data is to be kept safe and anonymous, and were permitted to withdraw from the study at any point.

### Subjects

Fourteen dogs were measured in this study. The age, sex, breed, weight and height of the animals are reported in Table 1. To assess the physical condition, we used a body condition score (BCS) for dogs, as developed and validated by Laflamme (1997).

**Table 1.**
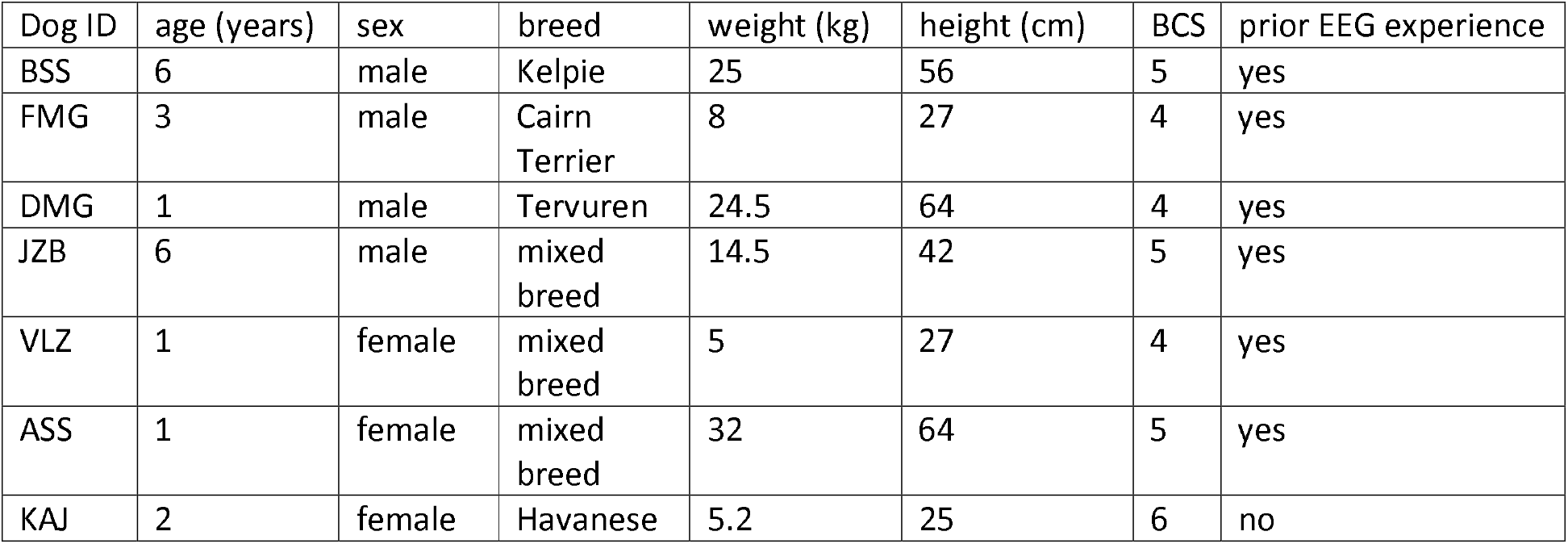
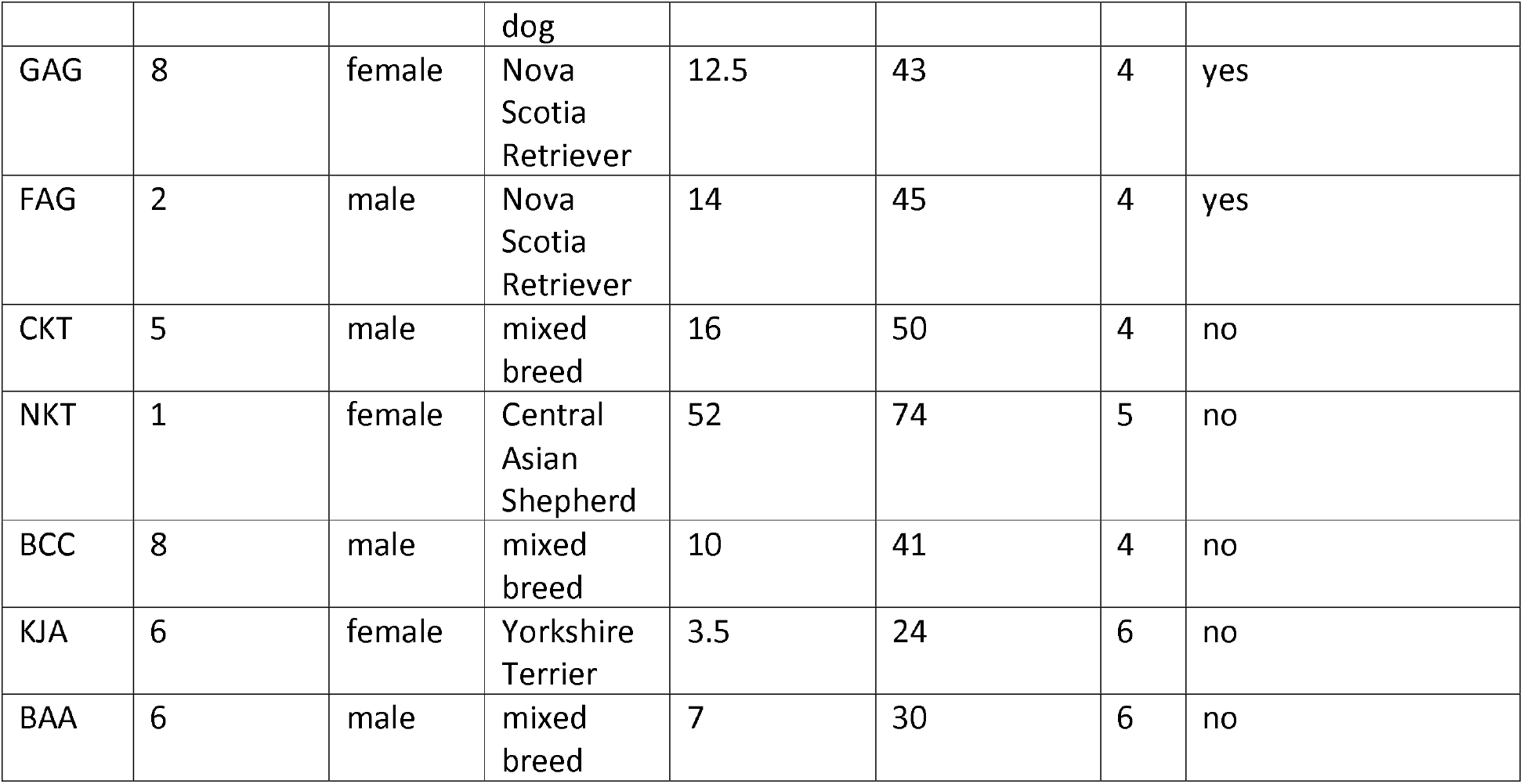
Identification codes and demographic data of the participating dogs.

### Behavioural paradigm

All dogs were allowed to explore the room and the experimenter prior to the experiment. Although the animals were thereby familiarized with the experimenter, most interactions, save for attaching the electrodes, were with the owner, and the dog was always physically closer to the owner as well. We did not tightly control how the owners managed their dog, but any peculiarities in the conduct of the dog or owner during the session were captured in a notebook and for twelve out of fourteen dogs, the sessions were also video-recorded.

For each dog, all recordings took place within the same experimental session to ensure that mostly identical factors characterize both experimental conditions: same room, same number of people present (owner and one experimenter), same time of day.

Two behavioural conditions were the source of our EEG recordings; they will be referred to below as ‘baseline’ and ‘test’. For the safety of our recording equipment and to minimize movement artefacts, it was necessary to somewhat restrict the animals’ movement in both conditions. Therefore, prior to the experiment, each dog was designated a spot on which to calmly reside. This might have required some self-control from the dogs (under the guidance of the owners), but it was a constant across both conditions. Only in the test condition was an additional delayed gratification challenge added; the dogs were awaiting permission to approach and i) eat a treat placed in front of them (N = 12) ii) take a favourite toy (N = 1) and iii) both (N = 1). The identity of the reward in this delayed gratification paradigm was deliberately allowed to vary between and sometimes within dogs, as a control against capturing activity that is specific to the identity of the stimulus or the stimulus category used in the tests. Another reason was to make sure that every dog was challenged with a treat of their liking, since in humans, inhibitory control in the absence of emotionally salient stimuli requires less theta band elevation (Andreu et al., 2019). The owners chose (or approved) which rewards were valued by their dog, based on their experience with the animal. Two dogs were tested with a toy. For GAG a toy was used instead of an edible treat, while ASS was tested with both. However, for ASS, only two toy trials were recorded, since the patience of the animal with the experiment ended the session after this (ASS increasingly tried to stand up and walk away, and after losing one of the electrodes, we ended the session for the safety of the equipment). A single toy trial attempted in a third dog (KAJ), was later discarded, because the owner could not confirm that the toy, which was novel to the dog, was to its liking, nor was a valid preference test used to establish if it is. A stricter inclusion rule for toy trials is based on previous work indicating that dogs can be very picky about toys (Kubinyi et al., 2020), but the discarded trial is still part of the shared data. The types of treat used for each dog are summarized in Table 2.

**Table 2.** Category, type, and cross-trial consistency (i.e. was an identical treat used on all trials) for each dog in the experiment.

| Dog ID | category | type | same across trials |
| --- | --- | --- | --- |
| BSS | edible | ham | yes |
| FMG | edible | sausage | yes |
| DMG | edible | sausage | yes |
| JZB | edible | special snack | yes |
| VLZ | edible | special snack | yes |
| ASS | edible, toy | special snack, ball | no |
| KAJ | edible | special snack | no |
| GAG | toy | stuffed animal | yes |
| FAG | edible | special snack | yes |
| CKT | edible | special snack | yes |
| NKT | edible | special snack | yes |
| BCC | edible | ham | yes |
| KJA | edible | special snack | no |
| BAA | edible | special snack | no |

EEG recordings in the test condition started when the forbidden treat was placed in front of the dog, within its reach, followed by a command, always issued by the owner, which instructed the animal to wait. After a ∼30-second-long EEG recording, the animal was allowed to eat the treat to remain motivated for further trials. Recording EEG on test trials was halted before the dog was allowed to approach the reward. Baseline recordings were randomly taken before, in between, and after test trials. To match the length of the recordings from each condition, we aimed for about 30 seconds duration in both, but report minor variation in the final data (standard deviation of 3 seconds). To be able to test if an effect is also visible on the individual level, we aimed to meet and exceed a minimum of ∼6 trials per condition in most dogs, following a recommendation by Camerlink and Pongrácz (2021). This could not be achieved in only two dogs due to the tight schedule of the testing room that day. The exact number of trials/condition and the randomized sequence in which they were recorded is displayed for each dog in Table 3.

**Table 3.** Sequence, number, and type of trials (T = test, B = baseline) for each dog.

| Dog ID | 1 | 2 | 3 | 4 | 5 | 6 | 7 | 8 | 9 | 10 | 11 | 12 | 13 | 14 | 15 | 16 | 17 | 18 | 19 | 20 | 21 |
| --- | --- | --- | --- | --- | --- | --- | --- | --- | --- | --- | --- | --- | --- | --- | --- | --- | --- | --- | --- | --- | --- |
| BSS | B | T | B | T | B | B | T | B | T | B | T | T | B |  |  |  |  |  |  |  |  |
| FMG | T | B | T | T | B | B | T | B | T | T | B | T |  |  |  |  |  |  |  |  |  |
| DMG | B | T | B | T | T | B | T | B | T | T |  |  |  |  |  |  |  |  |  |  |  |
| JZB | B | B | B | T | T | B | B | T | B | T | B | T | T | T | B | B |  |  |  |  |  |
| VLZ | B | B | T | B | T | T | B | B | B | T | T | B | B | T | B | T | B | T | B |  |  |
| ASS | T | T | B | B | B | T | T | B | B | T | T | T | B | B | B | B | T | T | B | T | B |
| KAJ | T | T | B | B | B | T | B | B | B | T | T | B | B | T | B | B | T |  |  |  |  |
| GAG | T | T | B | B | B | T | T | B | B | T | T | B | B | T | T | B | B |  |  |  |  |
| FAG | B | B | B | B | T | B | T | B | T | T | B | B | T | T | T | B | B |  |  |  |  |
| CKT | B | B | T | T | B | B | T | T | B | B | T | T | T | B | B | B | T | T |  |  |  |
| NKT | B | B | T | T | T | B | B | B | B | T | T | T | T | B | B | T | T |  |  |  |  |
| BCC | B | B | B | B | T | T | B | B | T | T | T | B | B | T | T |  |  |  |  |  |  |
| KJA | B | B | B | B | B | T | T | B | B | T | T | B | B | T | T | T |  |  |  |  |  |
| BAA | B | B | B | T | T | B | B | T | T | B | B | B | T | T | B | B | T | T |  |  |  |

### EEG recordings

The set-up used here to record EEG in awake dogs (Figure 1) is a variation on the one developed for sleep research in this species by Kis et al. (2014). Six gold-coated Ag/AgCl electrodes were attached for the duration of the experiment to the scalp of the dog with EC2 gel (Grass Technologies, USA), keeping the impedance below 15 kΩ. A 30-channel Flat Style SLEEP La Mont Headbox and an HBX32-SLP 32-channel preamplifier (La Mont Medical Inc., USA) were used to collect and pre-process the signals. The occipital bone was used to place a reference electrode, and the musculus temporalis for the ground electrode. Signals of interest were collected from the skull midline, with a frontally placed electrode (Fz), between the eyes, and a central electrode (Cz) between Fz and reference. To monitor eye movements, an additional two electrodes were placed on the zygomiotica (F7 and F8).

**Figure 1.**
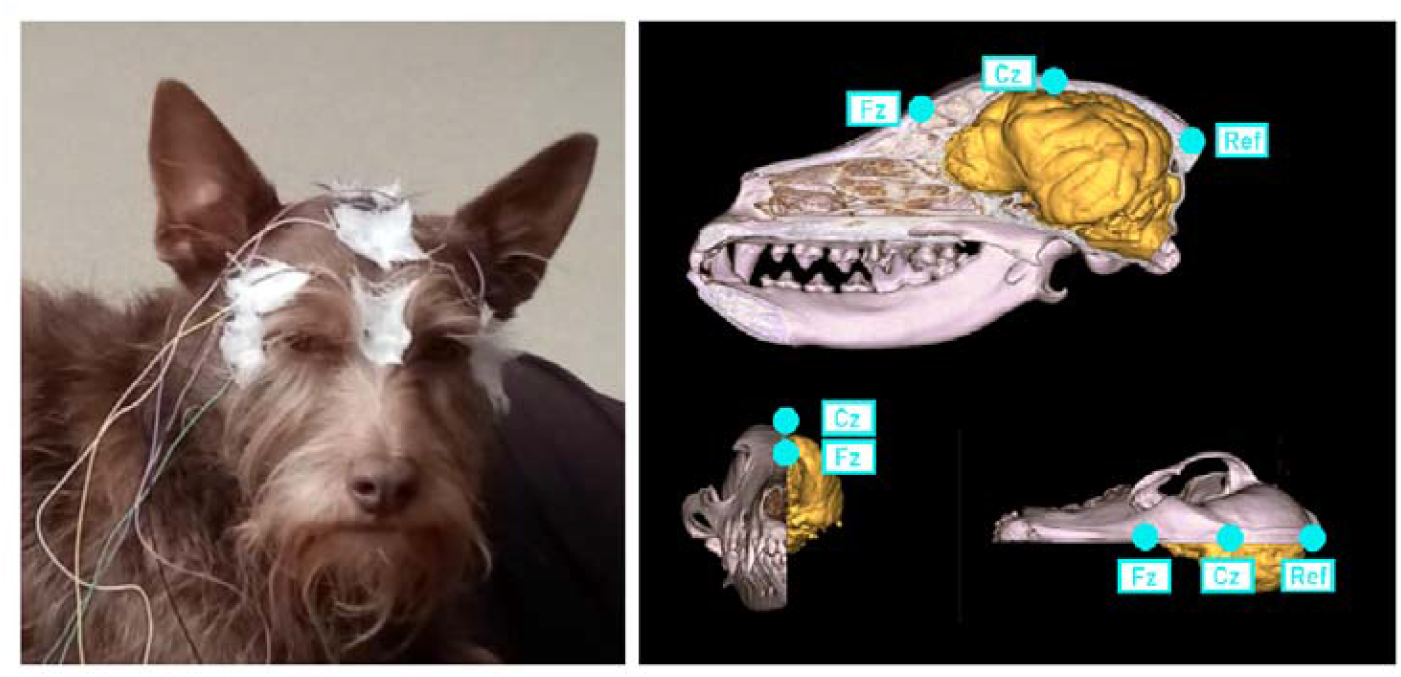
Placement of electrodes, real-life example (left side) and a 3D model of electrode position relative to the brain of a typical dog, highlighting, in particular, electrodes Fz, Cz, and reference (right side). 3D model by Kálmán Czeibert, used with permission.

### EEG post-processing

All filtering of the EEG prior analysis was done with second-order section Butterworth filters, similar to those in our previous work on dogs’ sleep EEG (Iotchev et al., 2023, 2019). The low-pass boundary was set to 30 Hz (passband) and 35 Hz (stopband). The high-pass boundaries were set to 0.1 Hz (passband) and 0.05 Hz (stopband). The 0.1 to 30 Hz passband range was used here to align with several previous works in humans (Batterink and Paller, 2017; Batterink and Zhang, 2022; Lyamin et al., 2008; Moser et al., 2021; Waterman et al., 1993) and our own previous analyses in dogs (Iotchev et al., 2024, 2023). Prior to filtering the signal, the EEG traces were cleaned for eye-movement artefacts. We applied a regression-based approach (Ronca et al., 2024), using the eye-movement tracking electrodes (F7 and F8) as predictors for either Fz or Cz activity, subtracting the explained variance from the raw EEG signal of each midline electrode. To minimize the loss of real signal that may be shared between the midline and eye-movement tracking electrodes, the latter were first low-pass filtered at a passband of 15 Hz (stopband 17 Hz), since the main spectral content of eye-movement artefacts was found to usually not exceed 15 Hz (Ronca et al., 2024).

### Frequency-power extraction

Frequency power extraction is closely modelled on our previous approach to quantifying spectral power in dogs, as a fraction of the total measured spectrum (Iotchev et al., 2023; Kis et al., 2017). We will refer to this measure henceforth as *relative theta power*.

Time-frequency analysis (*spectrogram*.*m* in Matlab) was applied across a shifting, 50% overlapping time-window of 4-second duration, zero-padded for a 0.1 Hz resolution. Theta was defined in the 5-7 Hz range (Jadeja, 2021) to minimize overlap with neighbouring frequency bands, i.e. delta and alpha. Generally, for our calculation of relative power in the bands alpha, beta, delta, and theta, all boundaries were adjusted to minimize overlap, which resulted in other minor differences from our previous work. Specifically, the lower boundary of beta was also moved up from 12 to 13 Hz to avoid overlap with alpha. The *relative theta power* calculation mostly followed our previous protocol for calculating relative power, except that medians instead of averages were used to integrate power, since this makes the estimates more robust to artefacts that may have been missed by the automatic cleaning. The spectrum within each band (alpha 8-12 Hz, beta 13-30 Hz, delta 1-4 Hz, theta 5-7 Hz) was first integrated across time-windows and then across frequencies within that band to end up with a single value for each frequency band within a recording. The power value for each band was then re-calculated as a fraction of the sum of the values for all four bands.

### Data and analyses

The final data comprised 226 recordings from 14 dogs. Of these, 120 were baseline recordings. As intended, most recordings were half a minute long: the mode and median of the distribution were at exactly 30 seconds. The range spanned from 18 to 48 seconds, therefore control analyses were conducted by checking for correlations between trial duration and relative theta power.

For our main analyses, since relative theta power, initially calculated as a percentage, can also be represented as part of a whole, we used Beta regression (with default link function), recommended for values limited between 0 and 1 (Geissinger et al., 2022) in R (R Development Core Team, 2021) to analyse condition differences. To account for which observations come from the same dogs, we applied Beta regression as a generalized linear mixed model (glmm), which was possible with the glmmTMB package (Brooks et al., 2017; McGillycuddy et al., 2025). The ID of the animals was used as a random factor. The same analysis without a random factor was used to test condition effects within each dog separately. For individual tests, we also calculated the combined probability (Fisher’s method (Fisher, 1970), applied through the sumlog function in R (Dewey, 2019)) to test if significant findings were rather hits or false positives. The former was deemed more likely if the combined probability of all individual-level tests was also significant.

## Results

### Control analyses

Trial duration was not correlated with relative theta power on Fz (P = 0.310) or Cz (P = 0.078).

### Behaviour

The dogs mostly succeeded in following the instructions of the test condition, and wait for up to 30 seconds until permission was given; sufficient to obtain a minimum of 6 valid test trials in each animal. Twice, a reward was prematurely consumed within the first few seconds (both times by the same dog, ASS). These trials were discarded before analysis.

### Group-level analyses

Analysing across all available recordings, accounting for which recordings come from the same dogs (random factor = dog ID), there was a significant condition effect (test > baseline) for relative theta power on Fz (z = 4.144, 95% CI: 0.10 - 0.29, P < 0.001), but not on Cz (P = 0.958). The distribution of relative theta power values is visualized in Figure 2A for both conditions.

**Figure 2.**
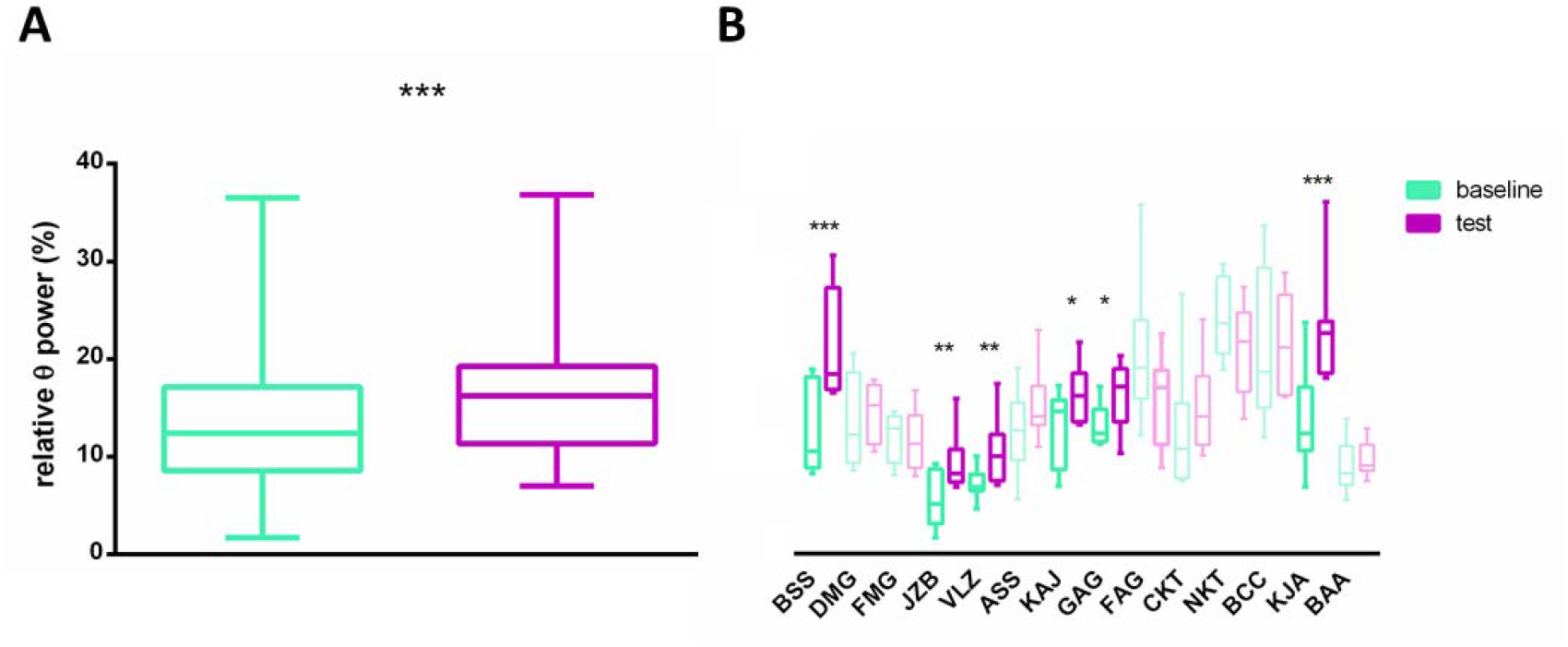
Data distribution for recordings observed on Fz, separated by condition (A), and for each dog (B). The data of dogs for which the effect was significant is plotted with stronger colour contrast. Significance levels are indicated with * (P < 0.05), ** (P < 0.01), and *** (P < 0.001).

### Individual analyses

Of the twelve dogs for which sufficient data were available for statistical analysis based on the ≥ 6 observations/condition cut-off criterion (Camerlink and Pongrácz, 2021), a condition effect for theta power (test > baseline) was significant in six. BSS (z = 3.503, P < 0.001), JZB (z = 3.154, P = 0.002), VLZ (z = 3.087, P < 0.002), KAJ (z = 2.410, P = 0.016), GAG (z = 2.323, P = 0.020), and KJA (z = 3.811, P < 0.001). In addition, a trend effect in the same direction was observed in ASS (P = 0.052), while in the remaining animals the effect was not significant (P > 0.1). The combined probability for the effects observed on Fz was significant (Fisher’s method, ⍰^2^ = 96.5, P < 0.001). These results are visualized in Figure 2B. On Cz, only KJA displayed a significant effect in the expected direction (test > baseline, z = 2.050, P = 0.040), while an opposite effect (baseline > test) was observed in ASS (z = −2.001, P = 0.045) and NKT (z = −2.014, P = 0.044). In no other dog was there a condition difference in theta power on Cz (P > 0.1), nor was the combined probability for effects observed on Cz significant (Fisher’s method, P > 0.05).

## Discussion

In this work, we present the results of a first inquiry into the utility of EEG for studying cognitive control in dogs. Theta frequency power, which in humans peaks over the frontal midline during various types of cognitive control tasks (Eisma et al., 2021; Inanaga, 1998), was also elevated in our study when comparing test and baseline EEG recordings across and within a sample of family dogs.

A first question to address is how we can be certain to have observed a true analogue of human theta in the domestic dog. This is an important question because it is hard to vary cognitive control demands without also varying the presence/absence of a rewarding stimulus. In humans, frontal midline theta rises with mental effort (Inanaga, 1998), whereas simply registering the presence of reward is instead linked to changes in higher frequency bands, like beta and gamma (Marco-Pallares et al., 2015). Two arguments can be made as to why we succeeded in measuring activity analogous to human frontal midline theta. One argument is based on what we already know about how typical oscillations of interest are expressed in humans, dogs, and other animals. The other argument will concern the localization of the here-observed effects along the scalp midline.

Several commonly studied EEG phenomena have a very similar frequency signature between humans and dogs, even when they are known (or suspected) to vary between humans and other animals. Spike-wave discharges, for example, the EEG expressions of absence seizures, are well-known to display a higher frequency in WAG/Rij rats, although they provide an otherwise strong model for human spike-wave complexes (Coenen and van Luijtelaar, 2003). Sleep spindles, on the other hand, for several animal species are suspected to be of much lower frequency than in humans, based on observational data (Nicol et al., 2000; Voirin et al., 2014). In dogs, however, both spike-wave discharges and sleep spindles are of the same frequency as in humans (Iotchev and Kubinyi, 2021; Musteata et al., 2023; Poma et al., 2010; Wielaender et al., 2017) and so are the frequency profiles of the different vigilance states, i.e. wakefulness, REM and deep sleep (Kis et al., 2014). In summary, there is currently little to no indication to expect a different frequency definition for theta in the dog.

Evidence for a humanlike theta response also derives from the localization of the effects across the scalp midline. When the group was analysed as a whole, relative theta power displayed a significant condition effect only on Fz. This was in agreement with individual tests on data from this electrode, where more dogs consistently displayed a significant condition effect in the expected direction (test > baseline). The relevance of topography in dog EEG data is supported by previous work, showing that humanlike differences between frontal and central electrodes can be replicated in dogs (Iotchev et al., 2019), although only two midline electrodes are commonly used in this species (Kulgod et al., 2025).

While for some research questions, stimulus consistency is crucial, in our case, it was important to depart from this standard. Stimulus-consistent tests do not help differentiate stimulus-driven from task-driven changes in EEG power. We were, however, primarily interested in the effect of the task. Moreover, using the same bait across participants would have risked that the value of the delayed gratification would be too low for some of the subjects. If the stimulus is close to emotionally neutral, lower theta-power elevations during inhibitory control challenges could be expected (Andreu et al., 2019), which would in turn decrease the signal-to-noise ratio for our variables of interest. For this reason, owners were encouraged to choose the treat based on their knowledge of the dog’s preferences and were given some freedom in how to bait the test. Some owners used the same type and amount of treats across trials, while others varied their offer from within a collection of favoured treats (see Table 2). By design, we tested one dog only with its favourite toy (GAG). None of these differences precluded us from seeing significant condition effects (test > baseline) when looking at dogs separately. Since all testing situations involved withholding an appetitive response for ∼30 seconds, we deem it valid to treat test trials within and across dogs as cumulative.

What we can overall ascertain is that for the here studied sample of dogs, neural activity reminiscent of human frontal-midline theta in both frequency and localization along the scalp midline, was elevated specifically during a delayed gratification challenge. The coincidence of a humanlike theta elevation from baseline, with a condition that factually requires inhibitory control, suggests, by analogy, that this behavior in dogs is under prefrontal control (Zavala et al., 2018). This is demonstrated for almost half of the sample studied here. While smaller samples tend to be underpowered (Schmidt and Hunter, 2004), and we therefore counted on the effect not showing up in all dogs, a combined probability test was also applied to further inquire if significant results on the individual level were more likely to be hits or false positives. We found support for the former only on Fz. The agreement between group-level and individual results for Fz-recorded theta-power strengthens our arguments, since our observation cannot be reduced to a group-level abstraction, nor is a single outlier subject driving the group-level effect.

This is the second known to us study to link dog behavior suggestive of cognitive control to neural activity, which implicates the frontal lobes. Previous work by Cook, Spivak and Berns (2016) demonstrated frontal lobe activation with the more reliable source localization of the fMRI technique. However, meaningful follow-up work for their exciting result is inherently limited by the cost of implementing fMRI (Montgomery, 2023), and the limited space within the scanner allows for little more than passive perception experiments, which dominate the literature on dog fMRI (Andics et al., 2016; Berns et al., 2012; Gábor et al., 2020; Thompkins et al., 2021). Moving outside the scanner allowed us to test a situation more reminiscent of what the animals are likely to encounter in day-to-day situations: being forbidden to eat food off the floor or the kitchen table, or having to wait for permission from their owner before approaching or acting upon interesting stimuli. The use of a more natural setting also allows us to derive more realistic assumptions as to when the animals make use of their ‘higher’ faculties. It is clear from previous work that dogs’ use of inhibitory self-control is not on par with that of humans. Dogs tend to continue to approach a wrong target in two-choice tasks, even when a reduced latency towards the right choice betrays that they are learning the difference (Bolló et al., 2025; Piotti et al., 2018). On the other hand, our results suggest that common situations for dogs, like abstaining from approaching or grabbing what the owner has forbidden, are likely solved by the engagement of cognitive control. This is contrary to the claim that most of dogs’ apparently cognitive behaviours can be reduced to instances of conditioning alone (Wynne, 2016).

A few important caveats and limitations also need to be discussed, as well as how our analysis and arguments account for these problems. Dogs’ waking EEG is inherently more noisy than that of humans (Kis et al., 2014), and this has led to a relatively large portion of EEG studies in dogs to focus on sleep physiology (Kulgod et al., 2025), when the animals are relatively motionless (but see e.g. Bálint et al. (2022) for exceptions). We have undertaken three measures to reduce this problem. Our first ‘line of defence’ was to adopt a simple, but well-established protocol for eye-movement correction - regression-based cleaning (Ronca et al., 2024), which allowed us to keep most of the signal, unlike the more common epoch rejection we so far applied in our sleep studies. Second, the use of medians instead of averages for integrating spectral power across time-windows and frequency bins allowed us to obtain estimates that are less likely to be skewed by unique and unusual events, i.e. by artefacts which survived the initial cleaning. Finally, focusing on a question for which the topography of the response matters also makes it less likely to observe a false positive confirmation of our hypothesis, since in dogs, artefacts should affect both midline electrodes equally. For making valid comparisons between Fz and Cz frequency power changes, relative power needed to be used, since absolute power would almost certainly always be lower on Cz in all bands, as Cz is closer to the reference electrode.

Another difficult question is what should count as the ideal baseline against which to compare our test condition. A dog, which is too calm during baseline recordings, may display differences in frequency power due to differences in its vigilance state, e.g. drifting into a drowsy state when nothing is happening. We deemed this particularly likely for baselines recorded before the first test trial, since during this stage, there was no task for the animal to attend. On the other hand, baseline trials recorded between test trials may capture anticipation of the need for cognitive control if the dog has begun to anticipate the tests. This, by itself, can raise theta-power and reduce the difference between conditions (Roelofs et al., 2006; Tóth-Fáber and Kóbor, 2025). For us, it was therefore important to vary between subjects (by virtue of using random sequences) how many baselines are recorded before versus between test trials. Significant condition differences did not seem to depend on this difference, however, and condition effects were observed on the individual level, regardless of the proportion of pre-test baselines to baselines recorded in between tests (compare Table 3 and individual test results). Since theta power elevation scales with task difficulty and mental effort, there are few, if any, circumstances which could mask condition effects. The additional challenge that our delayed gratification task imposed on top of the baseline adds to whatever difficulty (or comfort) the animals may otherwise experience. Not surprisingly, therefore, we observed significant condition effects in dogs with very different conduct during the recordings (see Supplementary notes and videos (the latter shared on OSF), e.g. KAJ adopted a resting position during baselines, while KJA attempted to shake off the electrodes on several occasions). Another difference in conduct we deemed irrelevant to the research question concerns the test condition: since theta waves appear to reflect reactive control (Mendl et al., 2023), it was not important if the dog had to be reminded not to eat the treats during tests, or managed to abstain with a single command at the start. In both cases succeeding depended on reacting to cues signalling the need for control, whether the cues are internal (the dog remembers the initial command) or external (the dog is reminded to stop). Follow-up work with larger samples can be used to thoroughly analyse which non-task-related dynamics make resting state theta fluctuate. Our first inquiry suggests that the delayed gratification challenge creates a strong enough contrast with most other internal states the dogs may be undergoing in the absence of an explicit task.

The ability to measure condition-induced changes in frontal midline theta opens up exciting possibilities for studying higher cognition in dogs. Although dogs have been argued to be a cognitively advanced species on the basis of findings from neuroscience (Bailey and Pereira, 2018), the neuroscientific approach to studying higher cognitive function in the dog is yet to be properly established. Cognitive control in particular is still mostly studied only through behaviour (Olsen, 2018). In humans, frontal midline theta has been implicated in both voluntary control (van Schie et al., 2022; Yamanaka and Yamamoto, 2010) and emergent awareness of implicitly learned patterns (Lu et al., 2023). As a note of caution, the clinical phenomenon of utilization (Lhermitte, 1983) suggests that prefrontal inhibition, of which theta is a surface EEG correlate (Zavala et al., 2018) may also operate as a background process. It may therefore be too early to speculate what frontal midline theta can teach us about the level of awareness or cognitive engagement in dogs. On the other hand, it is worth emphasizing that we were not measuring spontaneously ongoing background inhibition with this set-up. Instead, our paradigm introduces a salient change in the need for inhibitory control, not least by using treats tailored to each dog’s preference.

On a more practical note, identifying neuro-cognitive mechanisms of self-control around food can have important implications for animal welfare as obesity among family-owned dogs is an increasing problem (Lund et al., 2006). Poor cognitive control is an important comorbidity of obesity (Guxens et al., 2009; Kamijo et al., 2014; Sellaro and Colzato, 2017), ADHD (de Zeeuw et al., 2012), and depression (Grahek et al., 2019) in humans. These are conditions also of interest to the veterinary practitioner (Kealy et al., 2002; Laflamme, 1997; Senay, 1966). Therefore, a more intensive research into dogs’ cognitive control capacities can have practical implications for the veterinary field, while also strengthening the dog as an emerging model for human ADHD (Bunford et al., 2019; Kovács et al., 2025; Vas et al., 2007) and depression (Senay, 1966).

Our findings help set the stage for studying canine cognitive control across different behavioural paradigms and neuro-imaging modalities, therein complementing the first steps taken by Cook et al. (2016) in the direction of establishing a canine cognitive neuroscience.

## Supporting information

Supplementary_ExperimenterNotes

## Acknowledgements

The authors would like to cordially thank Sára Sándor for advice and feedback. All owners and dogs for their participation in this study. Hein van Schie, since the idea for this study was born out of a previous collaboration. Boglárka Morvai for discussions on statistics, which shaped some of the choices taken herein. Kauê Costa and Gilles van Luijtelaar for the critical thought exchange on the meaning of the findings.

## Data availability

The data for this study is shared on Open Science Framework (OSF) under: https://osf.io/9efcs/overview?view_only=4c05719bb6234dd9aab91f420821f306

## Author contributions

IBI developed the hypothesis and protocol, conducted the experiments, collected and analysed the data, and wrote the initial draft. MG provided methodological feedback, commented and edited the draft. AK acquired funding, provided methodological feedback, supervised the research, commented and edited the draft.

## References

Andics, A., Gabor, A., Gacsi, M., Farago, T., Szabo, D., Miklosi, A., 2016. Neural mechanisms for lexical processing in dogs. Science (80-.). 353, 1030–1032. 10.1126/science.aaf3777

Andreu, C.I., Palacios, I., Moenne-Loccoz, C., López, V., Franken, I.H., Cosmelli, D., Slagter, H.A., 2019. Enhanced response inhibition and reduced midfrontal theta activity in experienced Vipassana meditators. Sci. Rep. 9, 13215.

Bailey, J., Pereira, S., 2018. Advances in neuroscience imply that harmful experiments in dogs are unethical. J. Med. Ethics 44, 47–52. 10.1136/medethics-2016-103630

Bálint, A., Eleőd, H., Magyari, L., Kis, A., Gácsi, M., 2022. Differences in dogs’ event-related potentials in response to human and dog vocal stimuli; a non-invasive study. R. Soc. Open Sci. 9, 211769.

Ball, T., Schreiber, A., Feige, B., Wagner, M., Lücking, C.H., Kristeva-Feige, R., 1999. The role of higher-order motor areas in voluntary movement as revealed by high-resolution EEG and fMRI. Neuroimage 10, 682–694. 10.1006/nimg.1999.0507

Batterink, L.J., Paller, K.A., 2017. Online neural monitoring of statistical learning. Cortex. 10.1016/j.cortex.2017.02.004

Batterink, L.J., Zhang, S., 2022. Simple statistical regularities presented during sleep are detected but not retained. Neuropsychologia. 10.1016/j.neuropsychologia.2021.108106

Berns, G.S., Brooks, A.M., Spivak, M., 2012. Functional MRI in Awake Unrestrained Dogs. PLoS One 7, e38027.

Bolló, H., Carreiro, C., Iotchev, I.B., Gombos, F., Gácsi, M., Topál, J., Kis, A., 2025. The Effect of Targeted Memory Reactivation on Dogs’ Visuospatial Memory. eNeuro 12.

Bray, E.E., MacLean, E.L., Hare, B.A., 2014. Context specificity of inhibitory control in dogs. Anim. Cogn. 17, 15–31.

Brooks, M.E., Kristensen, K., van Benthem, K.J., Magnusson, A., Berg, C.W., Nielsen, A., Skaug, H.J., Maechler, M., Bolker, B.M., 2017. glmmTMB Balances Speed and Flexibility Among Packages for Zero-inflated Generalized Linear Mixed Modeling. R J. 9, 378–400.

Brucks, D., Marshall-Pescini, S., Wallis, L.J., Huber, L., Range, F., 2017. Measures of Dogs’ Inhibitory Control Abilities Do Not Correlate across Tasks. Front. Psychol. 8, 849.

Bunford, N., Csibra, B., Peták, C., Ferdinandy, B., Miklósi, Á., Gácsi, M., 2019. Associations among behavioral inhibition and owner-rated attention, hyperactivity/impulsivity, and personality in the domestic dog (Canis familiaris). J. Comp. Psychol. 10.1037/com0000151

Camerlink, I., Pongrácz, P., 2021. Getting the statistics right for your manuscript. Appl. Anim. Behav. Sci. 10.1016/j.applanim.2021.105333

Coenen, A.M.L., van Luijtelaar, E.L.J.M., 2003. Genetic animal models for absence epilepsy: a review of the WAG/Rij strain of rats. Behav. Genet. 33, 635–655. 10.1023/A:1026179013847

Cook, P.F., Spivak, M., Berns, G., 2016. Neurobehavioral evidence for individual differences in canine cognitive control: an awake fMRI study. Anim. Cogn. 19, 867–878.

Czeibert, K., Andics, A., Petneházy, Ö., Kubinyi, E., 2019. A detailed canine brain label map for neuroimaging analysis. Biol. Futur. 10.1556/019.70.2019.14

de Zeeuw, P., Weusten, J., van Dijk, S., van Belle, J., Durston, S., 2012. Deficits in Cognitive Control, Timing and Reward Sensitivity Appear to be Dissociable in ADHD. PLoS One 7, e51416.

Dewey, M., 2019. Metap: meta-analysis of significance values//R package version 11.

Eisma, J., Rawls, E., Long, S., Mach, R., Lamm, C., 2021. Frontal midline theta differentiates separate cognitive control strategies while still generalizing the need for cognitive control. Sci. Rep. 11, 14641.

Fisher, R.A., 1970. Breakthroughs in statistics: Methodology and distribution., in: Statistical Methods for Research Workers. Springer, New York, pp. 66–70.

Gábor, A., Gácsi, M., Szabó, D., Miklósi, Á., Kubinyi, E., Andics, A., 2020. Multilevel fMRI adaptation for spoken word processing in the awake dog brain. Sci. Rep. 10.1038/s41598-020-68821-6

Geissinger, E.A., Khoo, C.L.L., Richmond, I.C., Faulkner, S.J.M., Schneider, D.C., 2022. A case for beta regression in the natural sciences. Ecosphere 13, e3940.

Grahek, I., Shenhav, A., Musslick, S., Krebs, R.M., Koster, E.H.W., 2019. Motivation and cognitive control in depression. Neurosci. Biobehav. Rev. 102, 371–381.

Guxens, M., Mendez, M.A., Julvez, J., Plana, E., Forns, J., Basagana, X., Torrent, M., Sunyer, J., 2009. Cognitive function and overweight in preschool children. Am. J. Epidemiol. 170, 438–446.

Hesslow, G., 1994. Inhibition of classically conditioned eyeblink responses by stimulation of the cerebellar cortex in the decerebrate cat. J. Physiol. 476, 245–256.

Inanaga, K., 1998. Frontal midline theta rhythm and mental activity. Psychiatry Clin. Neurosci. 52, 555–566.

Iotchev, I.B., Bognár, Z., Tóth, K., Reicher, V., Kis, A., Kubinyi, E., 2023. Sleep-physiological correlates of brachycephaly in dogs. Brain Struct. Funct. 228, 2125–2136.

Iotchev, I.B., Kis, A., Turcsán, B., Tejeda Fernández de Lara, D.R., Reicher, V., Kubinyi, E., 2019. Age-related differences and sexual dimorphism in canine sleep spindles. Sci. Rep. 9. 10.1038/s41598-019-46434-y

Iotchev, I.B., Kubinyi, E., 2021. Shared and unique features of mammalian sleep spindles – insights from new and old animal models. Biol. Rev. 10.1111/brv.12688

Iotchev, I.B., Szabó, D., Turcsán, B., Bognár, Z., Kubinyi, E., 2024. Sleep-spindles as a marker of attention and intelligence in dogs. Neuroimage 303, 120916.

Jadeja, N.M., 2021. How to Read an EEG. Cambridge University Press.

Jobert, M., Poiseau, E., Jähnig, P., Schulz, H., Kubicki, S., 1992. Topographical analysis of sleep spindle activity. Neuropsychobiology 26, 210–217. 10.1159/000118923

Kamijo, K., Pontifex, M.B., Khan, N.A., Raine, L.B., Scudder, M.R., Drollette, E.S., Evans, E.M., Castelli, D.M., Hillman, C.H., 2014. The Negative Association of Childhood Obesity to Cognitive Control of Action Monitoring. Cereb. Cortex 24, 654–662.

Kealy, R.D., Lawler, D.F., Ballam, J.M., Mantz, S.L., Biery, D.N., Greeley, E.H., Lust, G., Segre, M., Smith, G.K., Stowe, H.D., 2002. Effects of diet restriction on life span and age-related changes in dogs. J. Am. Vet. Med. Assoc. 220, 1315–1320.

Kis, A., Szakadát, S., Gácsi, M., Kovács, E., Simor, P., Török, C., Gombos, F., Bódizs, R., Topál, J., 2017. The interrelated effect of sleep and learning in dogs (Canis familiaris); an EEG and behavioural study. Sci. Rep. 7, 41873. 10.1038/srep41873

Kis, A., Szakadát, S., Kovács, E., Gácsi, M., Simor, P., Gombos, F., Topál, J., Miklósi, Á., Bódizs, R., 2014. Development of a non-invasive polysomnography technique for dogs (Canis familiaris). Physiol. Behav. 130, 149–156. 10.1016/j.physbeh.2014.04.004

Kovács, T., Szű cs, V., Gácsi, M., 2025. Self-control is associated with the interaction of ADHD-like traits and training level in dogs. Vet. J. 314, 106483.

Kubinyi, E., Szánthó, F., Gilmert, E., Iotchev, I.B., Miklósi, Á., 2020. Human Expressions of Object Preference Affect Dogs’ Perceptual Focus, but Not Their Action Choices. Front. Psychol. 10.3389/fpsyg.2020.588916

Kulgod, A., van der Linden, D., Franca, L.G.S., Jackson, M., Zamansky, A., 2025. Non-invasive canine electroencephalography (EEG): a systematic review. BMC Vet. Res. 21, 73.

Laflamme, D., 1997. Developmental and validation of a body condition score system for dogs. Canine Pract. 22, 10–15.

Lhermitte, F., 1983. “Utilization behaviour” and its relation to lesions of the frontal lobes. Brain 106, 237–255. 10.1093/brain/106.2.237

Lu, Y., Guo, X., Weng, X., Jiang, H., Yan, H., Shen, X., Feng, Z., Zhao, X., Li, L., Zheng, L., Liu, Z., Men, W., Gao, J.H., 2023. Theta Signal Transfer from Parietal to Prefrontal Cortex Ignites Conscious Awareness of Implicit Knowledge during Sequence Learning. J. Neurosci. 43, 6760–6778.

Lund, E.M., Armstrong, P.J., Kirk, C.A., Klausner, J.S., 2006. Prevalence and risk factors for obesity in adult dogs from private US veterinary practices. Int. J. Appl. Res. Vet. Med. 4, 177.

Lyamin, O.I., Lapierre, J.L., Kosenko, P.O., Mukhametov, L.M., Siegel, J.M., 2008. Electroencephalogram asymmetry and spectral power during sleep in the northern fur seal. J. Sleep Res. 10.1111/j.1365-2869.2008.00639.x

Marco-Pallares, J., Münte, T.F., Rodriguez-Fornells, A., 2015. The role of high-frequency oscillatory activity in reward processing and learning. Neurosci. Biobehav. Rev. 49, 1–7.

McGillycuddy, M., Warton, D.I., Bolker, B.M., 2025. Parsimoniously Fitting Large Multivariate Random Effects in glmmTMB. J. Stat. Softw. 112, 1–19.

Medenica, T., Bokulic, E., Sedmak, G., 2025. The subthalamic nucleus: From comparative neuroscience to the presumptive role in canine cognition, behavior, and behavioral disorders. J. Vet. Behav. 82, 75–88.

Mendl, J., Banerjee, S., Fischer, R., Dreisbach, G., Köster, M., 2023. Control in context: The theta rhythm provides evidence for reactive control but no evidence for proactive control. Psychophysiology 61, e14625.

Montgomery, R.M., 2023. Comparing EEG and fMRI in Psychiatry: A Comprehensive Guide to Their Strengths, Weaknesses, and Applications [WWW Document].

Moser, J., Batterink, L., Li Hegner, Y., Schleger, F., Braun, C., Paller, K.A., Preissl, H., 2021. Dynamics of nonlinguistic statistical learning: From neural entrainment to the emergence of explicit knowledge. Neuroimage. 10.1016/j.neuroimage.2021.118378

Musteata, M., Stefanescu, R., Borcea, D.G., Solcan, G., 2023. Very-Low-Frequency Spike– Wave Complex Partial Motor Seizure Mimicking Canine Idiopathic Head Tremor Syndrome in a Dog. Vet. Sci. 10, 472.

Nicol, S.C., Andersen, N.A., Phillips, N.H., Berger, R.J., 2000. The echidna manifests typical characteristics of rapid eye movement sleep. Neurosci. Lett. 283, 49–52. 10.1016/S0304-3940(00)00922-8

Olsen, M.R., 2022. An investigation of two ostensibly inhibitory control tasks used in canine cognition. Appl. Anim. Behav. Sci. 256, 105770.

Olsen, M.R., 2018. A case for methodological overhaul and increased study of executive function in the domestic dog (Canis lupus familiaris). Anim. Cogn. 21, 175–195.

Piotti, P., Szabó, D., Bognár, Z., Egerer, A., Hulsbosch, P., Carson, R.S., Kubinyi, E., 2018. Effect of age on discrimination learning, reversal learning, and cognitive bias in family dogs. Learn. Behav. 46, 537–553. 10.3758/s13420-018-0357-7

Poma, R., Ochi, A., Cortez, M.A., 2010. Absence seizures with myoclonic features in a juvenile Chihuahua dog. Epileptic Disord. 12, 138–141.

R Development Core Team, 2021. R: A Language and Environment for Statistical Computing. R Foundation for Statistical Computing Vienna Austria.

Roelofs, A., van Turenhout, M., Coles, M.G., 2006. Anterior cingulate cortex activity can be independent of response conflict in Stroop-like tasks. Proc. Natl. Acad. Sci. 103, 13884–13889.

Ronca, V., Capotorto, R., di Flumeri, G., Giorgi, A., Vozzi, A., Germano, D., di Virgilio, V., Borghini, G., Cartocci, G., Rossi, D., Inguscio, B.M.S., Babiloni, F., Aricó, P., 2024. Optimizing EEG Signal Integrity: A Comprehensive Guide to Ocular Artifact Correction. bioengineering 11, 1018.

Schmidt, F.L., Hunter, J.E., 2004. Methods of Meta-Analysis: Correcting Error and Bias in Research Findings, Methods of Meta-Analysis: Correcting Error and Bias in Research Findings. 10.4135/9781483398105

Sellaro, R., Colzato, L.S., 2017. High body mass index is associated with impaired cognitive control. Appetite 113, 301–309.

Senay, E.C., 1966. Toward an animal model of depression: A study of separation behavior in dogs. J. Psychiatr. Res. 4, 65–71.

Swick, D., Ashley, V., Turken, A.U., 2008. Left inferior frontal gyrus is critical for response inhibition. BMC Neurosci. 10.1186/1471-2202-9-102

Szabó, D., Janosov, M., Czeibert, K., Gácsi, M., Kubinyi, E., 2023. Central nodes of canine functional brain networks are concentrated in the cingulate gyrus. Brain Struct. Funct. 228, 831–843.

Taylor, Alex H., Bastos, A.P.M., Brown, R.L., Allen, C., 2022. The signature-testing approach to mapping biological and artificial intelligences. Trends Cogn. Sci. 26, 738–750.

Terrier, G., Gottesmann, C., 1978. Study of cortical spindles during sleep in the rat. Brain Res. Bull. 3, 701–706. 10.1016/0361-9230(78)90021-7

Thompkins, A.M., Lazarowski, L., Ramaiahgari, B., Gotoor, S.S.R., Waggoner, P., Denney, T.S., Deshpande, G., Katz, J.S., 2021. Dog–human social relationship: representation of human face familiarity and emotions in the dog brain. Anim. Cogn. 10.1007/s10071-021-01475-7

Tóth-Fáber, E., Kóbor, A., 2025. Alpha and theta activity during reward anticipation are modulated by implicit expectations about sequential risk. bioRxiv 2.

van Noordt, S., Heffer, T., Willoughby, T., 2022. A developmental examination of medial frontal theta dynamics and inhibitory control. Neuroimage 246, 118765.

van Schie, H.T., Iotchev, I.B., Compen, F.R., 2022. Free will strikes back: Steady-state movement-related cortical potentials are modulated by cognitive control. Conscious. Cogn. 104, 103382.

Vas, J., Topál, J., Péch, É., Miklósi, Á., 2007. Measuring attention deficit and activity in dogs: A new application and validation of a human ADHD questionnaire. Appl. Anim. Behav. Sci. 10.1016/j.applanim.2006.03.017

Voirin, B., Scriba, M.F., Martinez-Gonzalez, D., Vyssotski, A.L., Wikelski, M., Rattenborg, N.C., 2014. Ecology and Neurophysiology of Sleep in Two Wild Sloth Species. Sleep 37, 753–761. 10.5665/sleep.3584

Waterman, D., Elton, M., Hofman, W., Woestenburg, J.C., Kok, A., 1993. EEG spectral power analysis of phasic and tonic REM sleep In young and older male subjects. J. Sleep Res. 2, 21–27. 10.1111/j.1365-2869.1993.tb00056.x

Wielaender, F., James, F.M.K., Cortez, M.A., Kluger, G., Neßler, J.N., Tipold, A., Lohi, H., Fischer, A., 2017. Absence Seizures as a Feature of Juvenile Myoclonic Epilepsy in Rhodesian Ridgeback Dogs. J. Vet. Intern. Med. 32, 428–432.

Wynne, C.D.L., 2016. What Is Special About Dog Cognition? Curr. Dir. Psychol. Sci. 25, 345–350.

Yamanaka, K., Yamamoto, Y., 2010. Single-trial EEG Power and Phase Dynamics Associated with Voluntary Response Inhibition. J. Cogn. Neurosci. 22, 714–727.

Zavala, B., Jang, A., Trotta, M., Lungu, C.I., Brown, P., Zaghloul, K.A., 2018. Cognitive control involves theta power within trials and beta power across trials in the prefrontal-subthalamic network. Brain 141, 3361–3376.

