## Supplementary_ExperimenterNotes for "A human-like EEG signature of cognitive control in the domestic dog"

~~ Experimenter notes ~~

for BSS (recorded in 2025)

-look away strategy, but not necessary looking to the owner

-was given the same type of treat to wait for on each test trial, quantity might have varied

-2nd of first two baselines was a corrupt trial (electrode came off)

for DMG (recorded in 2025)

-was looking at owner while waiting for permission to eat

-FMG started whining latest during the middle of DMG's trials

-baseline1 and food5 were corrupt trials (food5 was initiated without bait and therefore terminated promptly)

for FMG (recorded in 2025)

-reference might have been slightly anterior to the occipital bone

-overall extremely calm

-video-recorded on 20.11.2025

Both dogs (DMG, FMG) had to be fed by the owner, they did not approach the forbidden treats themselves

for JZB (recorded in 2025)

-was looking at owner while waiting permission to eat

-was given the same type and quantity of treat to wait for on each test trial

-video-recorded on 27.11.2025

for VLZ (recorded in 2025)

-received additional calming during the first two baseline trials

-received double the bait in the 10th trial (test condition)

-was occasionally filmed with a phone, close-up, by her owner

-video-recorded on 03.12.2025

for ASS (recorded in 2025)

-2nd and 8th trial went wrong (both were food-bait trials), not used in analyses

-only dog so far that clearly tried to stand up when nothing was happening, whined during both tests and baselines

-last two baited trials used a toy instead of food, these trials and subsequent/inter-trial baselines are denoted such that on demand they can be analyzed separately

-video-recorded on 05.12.2025

for KAJ (recorded in 2025)

-the type of treat changed between baited trials

-on some baselines the dogs seemed particularly relaxed (head down), but also lied on the electrodes

-at the end one toy trial was also recorded, but on second thought it was discarded, since the toy preference of this dog is unknown and the toy was improvised from the lab's own inventory without proper preference test, nor could the owner confirm or deny how much the dog wanted the toy

-video-recorded on 09.12.2025

for GAG (recorded in 2026)

-video recording starts only after the first trial (by mistake)

-a toy was used as bait in all trials

-owner used petting as a strategy to keep the dog relatively calm during baselines, but the dog was overall rather calm

-the dog needed explicit command to take the toy, rather than an explicit command to not grab it

-as yet, the dog showed clear motivation to receive permission for the toy (e.g. whines, this is also captured in the video)

-the numbering of the trials (i.e. number-label given to the file prior recording) was messed up, but not the sequence (some numbers were skipped during naming)

-video-recorded on 20.01.2026

for FAG (recorded in 2026)

-video recording starts only after the second trial (by mistake)

-owner used petting as a strategy to keep the dog relatively calm during baselines, but the dog was overall rather calm

-had to be fed by owner (after permission was given), did not approach the treats by itself

-video-recorded on 20.01.2026

for CKT (recorded in 2026)

-exposed to another dogs whines in the background, but this was constant throughout both conditions

-owner petted the dog to calm it down after wire-up

-dog needed reminders to not eat the treat after initial command, for at least the first few trials (check video, too)

-during baseline recordings dog showed signs of relaxation (lied on its side)

-food trial 3 may be too short for analysis (headbox connection was lost after 14 seconds)

-video-recorded on 21.01.2026

for NKT (recorded in 2026)

-tried to stand up at least once in the beginning, from the middle until the end it seemed to stay calm in one place without explicit commands

-owner recorded the dog with their phone while the epxeriment was ongoing, but that was a constant across conditions

-video-recorded on 21.01.2026

for BCC (recorded in 2026)

-tried to remove the electrodes on at least 3 occasions (and during baseline recordings)

-one electrode had to be re-attached (twice) after trial 5 (first food trial)

-received calm touch to stay calm

-before trial 9 (food trial) the treat was brought closer in front of the dog

-at least once had to be sharply reminded to not eat the treat

-video-recorded on 28.01.2026

for KJA (recorded in 2026)

-electrode(s) came off between trial 2 and 3 (baselines), were re-attached prior recordings

-received calm speech and petting during most baseline trials, for the first two, three trials was sitting in the owner's lap

-tried to shake off the electrodes on multiple occasions

-was exposed to the whining of another dog (with whom she shares the household and who was waiting to be recorded later)

-during test trials the treat was placed extremely close to her

-had to be occasionally reminded to not eat the treat before permission was given

-maintained eye-contact with owner during delayed gratification waiting time

-video-recorded on 31.01.2026

for BAA (recorded in 2026)

-different electrodes came off on a few occasions, were re-attached in time (before any recording)

-on at least one occasion had to be reminded to not eat the forbidden treat

-maintained eye-contact with owner during test trials

-received calm speech and petting during most baseline trials

-on at least one occasion tried to run off between recordings, but was generally calm, possibly misunderstood being given a signal to run free

-occasionally exposed to whines from the other dog (KJ) but rather rarely (KJ remained mostly calm while waiting BA to finish)

-video-recorded on 31.01.2026
